# Ivermectin converts cold tumors hot and synergies with immune checkpoint blockade for treatment of breast cancer

**DOI:** 10.1101/2020.08.21.261511

**Authors:** Dobrin Draganov, Zhen Han, Nitasha Bennett, Darrell J. Irvine, Peter P. Lee

## Abstract

We show that treatment with the FDA-approved anti-parasitic drug ivermectin induces immunogenic cancer cell death (ICD) and robust T cell infiltration into breast tumors. As an allosteric modulator of the ATP/P2×4/P2×7 axis which operates in both cancer and immune cells, ivermectin also selectively targets immunosuppressive populations including myeloid cells and Tregs, resulting in enhanced Teff/Tregs ratio. While neither agent alone showed efficacy *in vivo*, combination therapy with ivermectin and checkpoint inhibitor anti-PD1 antibody achieved synergy in limiting tumor growth (p=0.03) and promoted complete responses (p<0.01), also leading to immunity against contralateral re-challenge with demonstrated anti-tumor immune responses. Going beyond primary tumors, this combination achieved significant reduction in relapse after neoadjuvant (p=0.03) and adjuvant treatment (p<0.001), and potential cures in metastatic disease (p<0.001). Statistical modeling confirmed bona fide synergistic activity in both the adjuvant (p=0.007) and metastatic settings (p<0.001). Ivermectin has dual immunomodulatory and ICD-inducing effects in breast cancer, converting ‘cold’ tumors ‘hot’, thus represents a rational mechanistic partner with checkpoint blockade.

---

Checkpoint blockade (1, 2) has emerged as a revolutionary approach that harnesses a patient’s own immune system to treat cancer. However, checkpoint inhibitors as single agents are only effective in a subset of patients and cancer types (2). Recent studies suggest that efficacy of checkpoint inhibitors is primarily limited to cancers already infiltrated by T cells – often termed ‘hot’ tumors. In contrast, ‘cold’ tumors have little to no T cell infiltration and generally do not respond to checkpoint blockade. Early clinical studies with checkpoint blockade therapy in breast cancer have focused on triple negative breast cancer (TNBC), because this subtype has a higher mutational load and is thought to be more ‘immunogenic’ (3). Even so, anti-PD1/PDL1 antibodies have produced clinical responses in only a small subset (15-20%) of TNBC patients (4). As such, there is considerable interest in identifying drugs capable of priming breast tumors (turning ‘cold’ tumors ‘hot’) to synergize with checkpoint blockade.

A recently described phenomenon, termed immunogenic cell death (ICD) (5, 6), is a form of cell death that induces an immune response from the host. ICD is distinguished from classical apoptosis and other non-immunogenic or tolerogenic forms of cell death by several hallmarks, including release of ATP and HMGB1 and surface exposure of calreticulin (5-7). In cancer patients, ICD-based anti-tumor immune responses are linked to beneficial outcomes produced by some conventional chemotherapeutic agents (8-11). For example, efficacy of anthracyclines in breast cancer (12-14) and oxaliplatin in colorectal cancer (15) correlates with post-treatment increases in the ratio of cytotoxic CD8+ T lymphocytes to FoxP3+ regulatory T cells within the tumor. In contrast, poor responses to chemotherapy in solid tumors are associated with lymphopenia (16). Thus, ICD-inducing chemotherapy appears to work in conjunction with the host immune system to achieve efficacy. However, chemotherapy is a double-edged sword: it can suppress as well as stimulate immune cells. An agent that induces ICD of cancer cells without suppressing immune function would be ideal for combination with checkpoint blockade. Seeking such an agent among FDA-approved drugs, our group found that the anti-parasitic agent ivermectin promotes ICD in breast cancer cells(17). Among our previous findings was evidence that ivermectin, an anti-parasitic drug used worldwide since 1975, modulates the P2×4/P2×7 purinergic pathway, suggesting that ivermectin may further harness tumors’ intrinsic high extracellular levels of ATP for anti-cancer activity. Of note, P2×4/P2×7 receptors are widely expressed on various immune subpopulations, suggesting that ivermectin might also have direct immunomodulatory effects.

## Results

### Ivermectin can turn ‘cold’ breast tumors ‘hot’

Motivated by these findings, we studied the effects of ivermectin *in vivo* using the 4T1 mouse model of TNBC. HMGB1 is a chromatin protein present in all cells and its release is a hallmark of ICD (18). HMGB1 staining (green) was observed uniformly across the entire tumor from untreated mice (Fig. 1A). In contrast, tumors isolated from mice treated with ivermectin showed large areas of DAPI-positive cells lacking HMGB1 (Fig. 1B), suggesting that HMGB1 had been released into the extracellular space. Ivermectin treatment also altered calreticulin expression, with higher levels (green) observed in tumors from treated animals, indicating a significant increase in this ICD-associated prophagocytic signal and mediator (Fig. 1C-D). Robust infiltration of both CD4^+^ and CD8^+^ T cells was seen in ivermectin-treated tumors (Fig. 1F) but not in untreated tumors (Fig. 1E). Significantly higher percentages of cells were positive for CD4 (p<0.01, Fig. 1G) and CD8 (p<0.0001, Fig. 1H) in ivermectin-treated than in untreated tumors. Together, these data indicate that treatment with ivermectin induced hallmarks of ICD within 4T1 breast tumors and recruited large numbers of CD4^+^ and CD8^+^ T cells into these tumors. To further confirm that ivermectin induces ICD *in vivo*, we also utilized a classical vaccination approach considered as the gold standard for detection of ICD: by treating 4T1 cells with IVM to induce ICD *in vitro* as well as to prevent tumor outgrowth after inoculation into naïve mice, followed by subsequent challenge with live 4T1 cells (18). This experiment validated the induction of *bona fide* ICD by demonstrating protection against subsequent challenge with live 4T1 cells (p<0.01, Fig. 1I).

**Fig 1.**
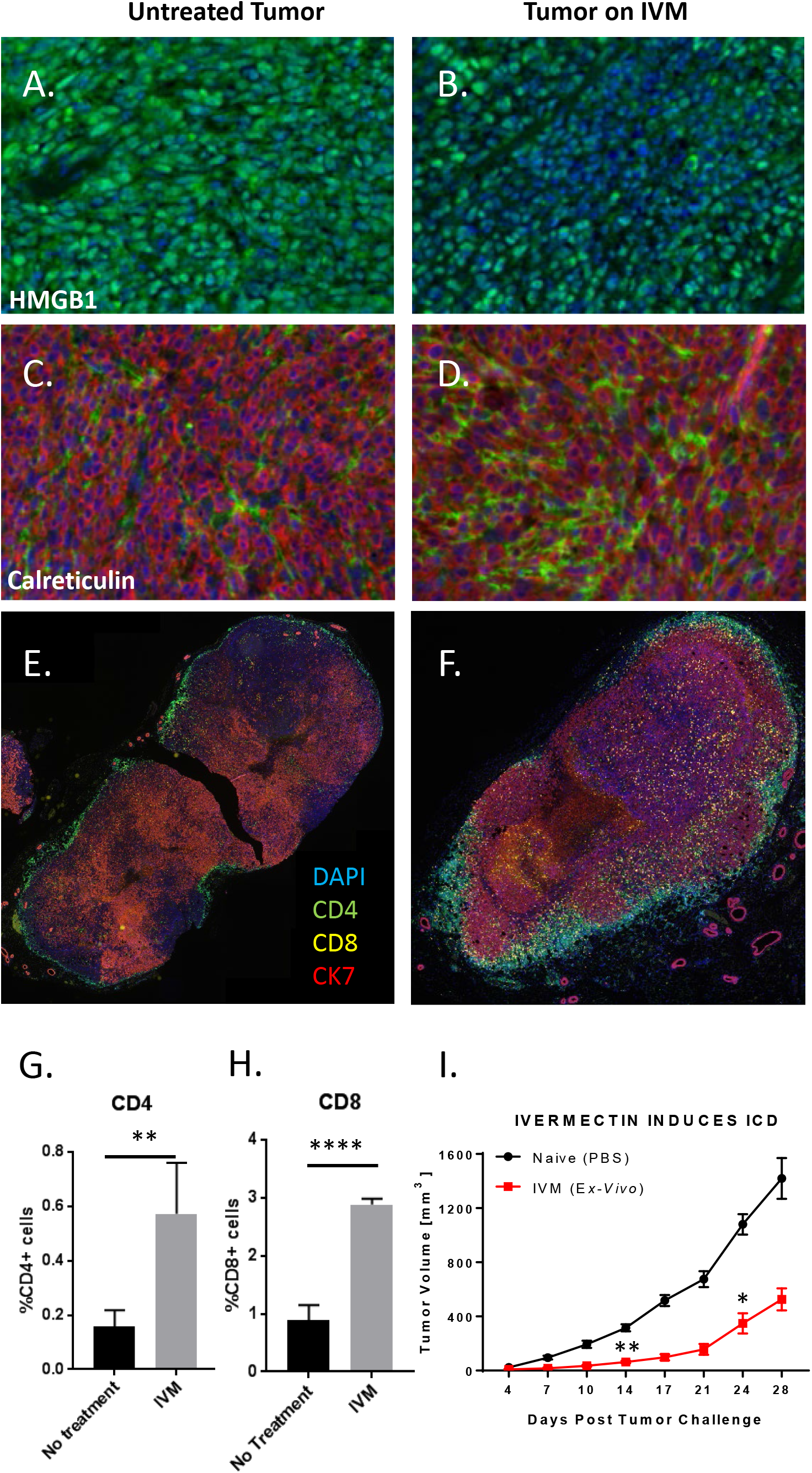
Treatment with ivermectin induces immunogenic cell death (ICD) in vivo and recruitment of T cells into tumors. 4T1 breast tumors were isolated from mice that were untreated (left panels) or ivermectin-treated (right panels) daily for 14 days. Figs. 1A, 1B show staining for HMGB1 (green), a hallmark of ICD. Figs. 1C, 1D show staining for calreticulin (green), another hallmark of ICD. Staining for CK7 (red) identifies 4T1 cells. Data are representative of two independent experiments. Figs. 1E, 1F show staining for CD4+ (green), CD8+ T cells (yellow), and cancer cells via staining for CK7 (red). Data are representative of three independent experiments. Figs. 1G and 1H display quantitative data on T cell infiltration shown in Figs 1E, 1F. Data were obtained by quantifying 5 random fields from whole tumor images. Fig. 1I demonstrates the protective effect of prophylactic subcutaneous vaccination with 1 million 4T1 cells treated with 12 μM ivermectin *ex vivo* (24h), then challenged contralaterally with live 4T1 cells one week post vaccination (n=4). Statistical significance was evaluated using the linear mixed effects model of log tumor volume. * p ≤ 0.05, ** p ≤ 0.01, *** p ≤ 0.001, **** p ≤ 0.0001.

### Direct immunomodulatory effects of Ivermectin

Ivermectin treatment *in vivo* did not produce any significant changes in the frequencies of various effector and regulatory CD4 (Fig. S2A) or CD8 (Fig. S2B) T cell subpopulations isolated from the spleens of treated animals. However, functional interrogation of splenocytes isolated from control vs. 4T1 tumor-bearing mice revealed significant immunomodulatory effects. Tumor-bearing mice one month post-inoculation developed enlarged spleens with an expanded population of CD11b+ myeloid cells (Fig. 2A). Ivermectin treatment *ex vivo* preferentially depleted this expanded CD11b+ myeloid population, normalizing the balance between myeloid and T cell compartments (Fig. 2A). Myeloid and lymphoid cell populations showed differential sensitivity to increasing doses of Ivermectin (Fig. 2B, S2C). A linear mixed effects model of log cell count adjusted for cell type revealed that CD11b+ myeloid cells were the most sensitive to ivermectin, showing significant reductions with as little as 4 μM after 48 hours, 8 μM after 24 hours, or 16 μM after 4 hours – demonstrating rapid and selective targeting of this immunosuppressive population (each result, p<0.0001). In contrast, achieving similar reductions in CD4 or CD8 T cells required higher doses and/or longer exposure to ivermectin: observed in CD8 T cells only after 48 hours of 8 μM or 24 hours of 16 μM, and in CD4 T cells only after the maximum exposure (48 hours of 16 μM). Consistent with ivermectin being an allosteric modulator of the ATP/P2×4/P2×7 signaling axis which operates in both cancer and immune cells, differential sensitivity in myeloid cells was P2×7-dependent (Fig. 2C). P2×7 blockade with 10 μM KN62 reversed the *ex vivo* depletion of CD11b+GR-1+ myeloid-derived suppressor cells (MDSC), CD11b+GR-1-Monocytes/Macrophages (Mon/Mac), and other immune subsets by ivermectin (p<0.001). To mimic more physiologically relevant conditions of exposure, we also treated splenocytes with lower non-cytotoxic doses of ivermectin and observed that over extended exposure, ivermectin had a significant potentiating effect on PHA-stimulated T cells and augmented the ratios of both CD8+ and CD4+ Teff/Tregs (Fig. 2D). The immuno-potentiating effects of extended exposure to lower non-cytotoxic doses of ivermectin was enhanced upon TCR stimulation (via PHA) and was inhibited in splenocytes from tumor-bearing mice (Fig. 2D), where different mechanisms including MDSCs as well as PD-1-mediated immunosuppression are known to interfere with proper TCR signaling and function.

**Fig 2.**
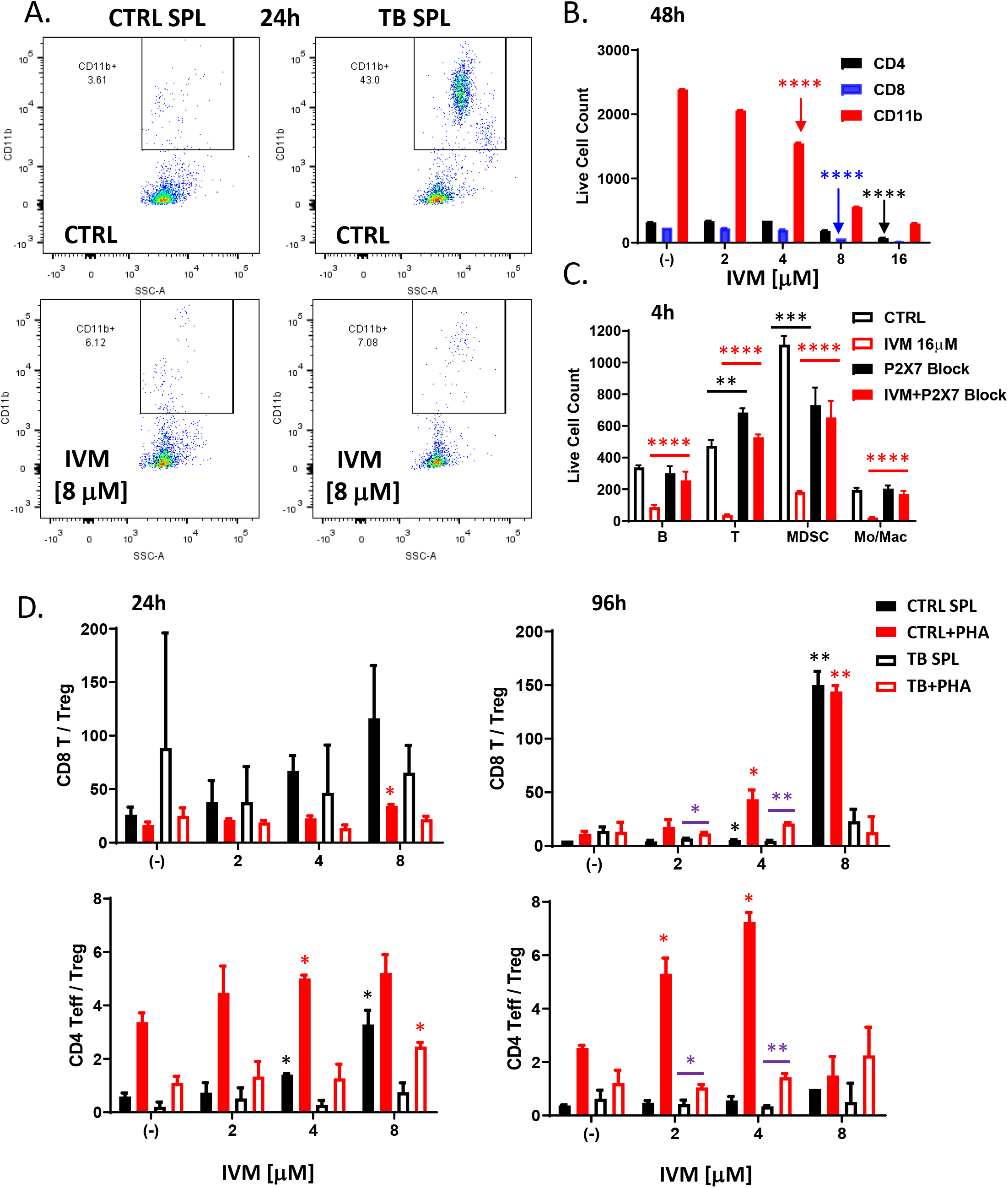
Immunomodulatory effects of ivermectin *ex vivo*. Splenocytes were isolated from the spleens of control naïve mice (CTRL) or untreated 4T1 tumor bearing mice (TB), one month post tumor inoculation, then cultured on 96-well tissue culture-treated plates in complete R10 medium for 4h-48h and analyzed by flow cytometry for spontaneous and ivermectin-induced changes in various immune subpopulations. (A) Depletion of the expanded CD11b+ myeloid cells isolated from the spleens of tumor-bearing mice by ivermectin treatment ex vivo. (B, C) Splenocytes isolated from 4T1 tumor-bearing mice were exposed to increasing doses of ivermectin for 4h or 48h showing differential dose- and time-dependent sensitivity of different immune subpopulations (see also Fig. S2C). Depletion of CD11b+GR-1+ MDSCs, CD11b+GR-1-Monocytes/Macrophages, CD19+ B cells and CD3+ T cells by IVM could be reversed by an inhibitor of P2×7/CaMKII (KN62 at 10 μM). (D) Splenocytes from naïve/untreated (CTRL) and 4T1 tumor-bearing (TB) mice were incubated for 24h and 4 days with increasing doses of Ivermectin (1-16 μM) with or without PHA to mimic TCR stimulation. Plots show averages and standard deviation based on triplicates; data representative of at least two independent experiments. Statistical significance versus (-) CTRL or as indicated was evaluated using the linear mixed effects model of log cell count adjusted for cell type: * p ≤ 0.05, ** p ≤ 0.01, *** p ≤ 0.001, **** p ≤ 0.0001.

### Ivermectin synergizes with anti-PD1 antibody to control tumor growth and induces protective immunity

The anti-cancer ICD and direct immunomodulatory effects of ivermectin raised the possibility that it could be combined with checkpoint blockade. We next investigated the efficacy of ivermectin and anti-PD1 antibody, alone or in combination, relative to no treatment (schema in Fig. S1A). Mean tumor volume over time was significantly decreased by the ivermectin and anti-PD1 antibody combination relative to no treatment (p<0.001, Fig. 3A). Through a joint statistical model of longitudinal tumor volumes, ivermectin and anti-PD1 antibody demonstrated synergistic activity, defined as an effect that is significantly greater than the sum of the drugs’ individual effects (submodel p=0.008, false discovery rate/FDR 3%, Table 1A). Complete tumor resolution was observed in 6/15 mice on the combination treatment, 1/20 on ivermectin alone, 1/10 on anti-PD1 antibody alone, and 0/25 on no treatment. Mice that resolved tumors on the ivermectin and anti-PD1 combination therapy were re-challenged with 100,000 4T1 cells in the contralateral mammary fat pads. All of these mice resisted development of new tumors (Fig. 3B), while control naïve animals all developed tumors (data not shown). This suggests that combined treatment with ivermectin and anti-PD1 induces protective anti-tumor immunity in complete responders.

**Table 1.** 1A. Tumor Growth in Primary Treatment 1B. Relapse-Free Survival in Adjuvant Setting 1C. Relapse-Free Survival in Metastatic Setting

| Tumor Growth per (log)Day | Estimate (SE) | Effect of Treatment (SE) | Submodel p value | False Discovery Rate |
| --- | --- | --- | --- | --- |
| No Treatment | 3.47 (0.18) |  |  |  |
| A – Ivermectin only |  | -0.84 (0.26) |  |  |
| B – anti-PD1 only |  | -1.46 (0.30) |  |  |
| Beyond Product of A + B |  | -1.05 (0.38) | 0.008 | 3% |

**1B. Relapse-Free Survival in Adjuvant Setting**
| Parameter | Hazards Ratio (95% CI) | Submodel p value | False Discovery Rate |
| --- | --- | --- | --- |
| No Treatment | 1.00 |  |  |
| A – Ivermectin only | 0.91 (0.36-2.26) |  |  |
| B – anti-PD1 only | 0.84 (0.33-2.10) |  |  |
| Beyond Product of A + B | 0.12 (0.03-0.54) | 0.007 | 2% |
| Adding IL-2 to A + B | 0.68 (0.22-2.13) | 0.51 | (67%) |

**1C. Relapse-Free Survival in Metastatic Setting**
| Parameter | Hazards Ratio (95% CI) | Submodel p value | False Discovery Rate |
| --- | --- | --- | --- |
| No Treatment | 1.00 |  |  |
| A – Ivermectin only | 1.48 (0.41-5.39) |  |  |
| B – anti-PD1 only | 0.75 (0.21-2.72) |  |  |
| Beyond Product of A + B | 0.02 (0-0.16) | <0.001 | <1% |
| Adding IL-2 to A + B | 0.58 (0.05-6.18) | 0.64 | (73%) |

**Fig 3.**
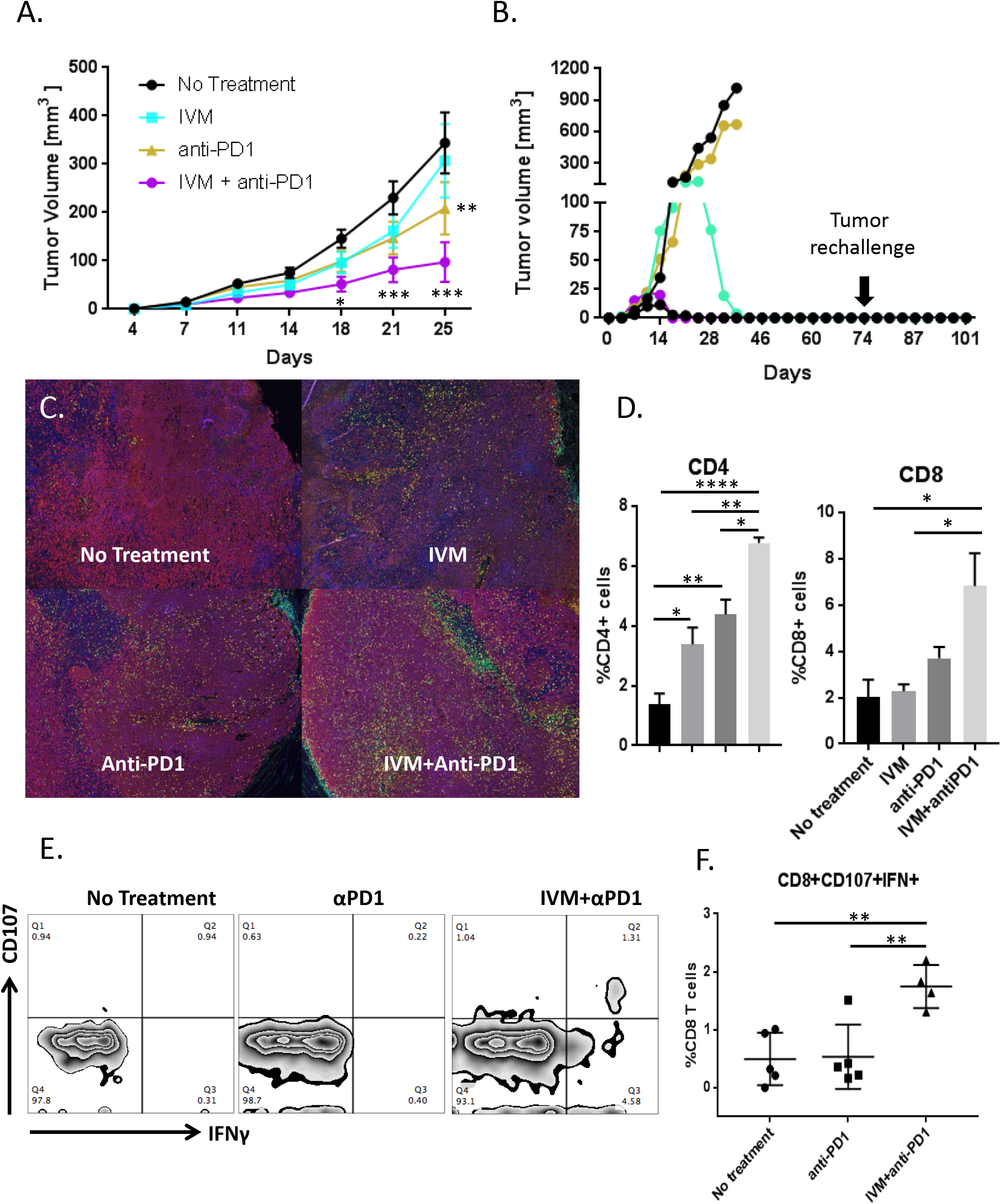
Ivermectin synergizes with anti-PD1 therapy to control tumor growth in vivo. Mice were inoculated with 100,000 4T1 cells four days before initiating therapy with ivermectin alone (n=20), anti-PD1 antibody alone (n=10), both drugs (n=15), or no treatment (n=25). **(A)** Tumor volume in control and treated animals. * p ≤ 0.05. ** p ≤ 0.01. *** p ≤ 0.001. **(B)** Tumor growth in individual animals treated with ivermectin plus anti-PD1 antibody (5 individual mice from one representative of three experiments shown). Three of five combination treated animals completely resolved their tumors. Animals that resolved tumors were re-challenged with 100,000 4T1 cells on the contralateral mammary fat pad 30 days after the termination of therapy. Mice were observed and palpated twice a week for an additional 30 days for the establishment of a tumor mass. **(C-F)** Combination therapy with ivermectin and anti-PD1 recruits significantly more T cells into tumor sites and generates tumor-reactive CD8+ T cells. Tumors were isolated from mice at Day 21. Staining was performed for nuclei (blue), CD4+ (green) cells, CD8+ cells (yellow), and tumor cells (red) **(C)**. Percent positive for CD4 or CD8 was measured in 5 random fields in each group and divided by the number of nuclei in the field **(D)**. Data are representative of two independent experiments. * p ≤ 0.05, ** p ≤ 0.01, **** p ≤ 0.0001. Splenocytes isolated from tumor-bearing mice that received no treatment (n=5), anti-PD1 alone (n=5), or ivermectin with anti-PD1 (n=4) were co-cultured with 4T1 cells. Reactive CD8+ cells were determined by CD107 mobilization and expression of IFNγ by flow cytometry. Representative flow plots for each treatment group are shown in **(E). (F)** percentage of CD8+ T cells reactive against 4T1 per mouse, grouped by treatment. ** p ≤ 0.01.

To gain further insight into the mechanism underlying efficacy of the combined treatment, we compared the magnitude to which ivermectin, anti-PD1, and their combination potentiated the infiltration of T cells. As shown visually in Fig. 3C and quantitatively in Fig. 3D, infiltration of both CD4^+^ and CD8^+^ T cells into 4T1 tumors (Day 21) was greatest after treatment with the combination of ivermectin and anti-PD1. To measure anti-tumor T cells, splenocytes were isolated from untreated, single agent treated or ivermectin plus anti-PD1 combination treated mice, then co-cultured with 4T1 cells as targets to measure CD107 mobilization and IFN-γ expression as markers for functional T cell responses (19). A functional tumor-specific immune response was confirmed by the presence of a discrete population of CD8^+^ T cells positive for CD107 and IFN-γ in mice treated with ivermectin plus anti-PD1, but not in mice treated with anti-PD1 alone or untreated controls (p<0.01; Fig. 3E, F).

### Combination therapy effective across spectrum of clinically relevant settings

Moving beyond control of primary tumors, we sought to test this combination immunotherapy across the major clinically relevant settings: neoadjuvant, adjuvant, and metastatic treatments. We also explored the effects of further augmenting this combination immunotherapy with Interleukin-2 (IL-2). IL-2 was the first cytokine to be successfully used in the treatment of cancer to induce T cell activation(20). A major challenge in the development of IL-2 as a therapeutic antitumor agent is that IL-2 can act on both T cells and regulatory T cells (Tregs). The contrasting actions of IL-2 has led to inconsistent responses and limited the development of high-dose IL-2 for cancer immunotherapy. Increasing the half-life of IL-2 has been shown to be a promising strategy for improving IL-2 based immunotherapy. This can greatly reduce the dose of IL-2 required for therapeutic activity, enhancing both safety and efficacy (21, 22). We explored the secondary hypothesis that addition of a recombinant albumin-IL-2 fusion with extended half-life to the ivermectin and anti-PD1 regimen (anti-PD-1 + IL-2 therapy, termed “IP” for simplicity) can further improve the efficacy of our combined treatment.

Neoadjuvant therapy has come to play an increasingly prominent role in the treatment of cancer. We tested treatment of ivermectin combined with anti-PD-1 and IL-2 (IP) by monitoring survival of animals receiving neoadjuvant combination therapy followed by surgical resection of the primary tumor on day 16 following tumor inoculation (schema in Fig. S1A). Development of loco-regional recurrence and distant metastases were monitored by bioluminescent imaging, and animals were euthanized upon decline in body condition score and signs of morbidity. All untreated animals required euthanasia due to lethal diseases around day 20-25 following surgical resection of primary tumor (Fig. 4A). Treatment with IP therapy alone provided some survival benefit with approximately 40% of animals remaining free of lethal disease. The best survival outcome was seen with the combination of IP and ivermectin therapy, with approximately 75% of animals becoming long-term survivors following surgical resection (p<0.05, Fig. 4A). Surviving treated mice were re-challenged with 100,000 4T1 cells in the contralateral mammary fat pads. The majority of IVM + IP treated mice did not develop new tumors (Fig. 4B), while IP treated and control naïve nice all developed tumors. Splenocytes from these animals were reactive (via ELISPOT) against 4T1 cells, demonstrating evidence for anti-tumor T cell responses in the IVM + IP treated animals (Fig. 4C). These results suggest that the IVM + IP combination treatment is effective in the neoadjuvant setting and induces protective anti-tumor immunity in responders.

**Fig 4.**
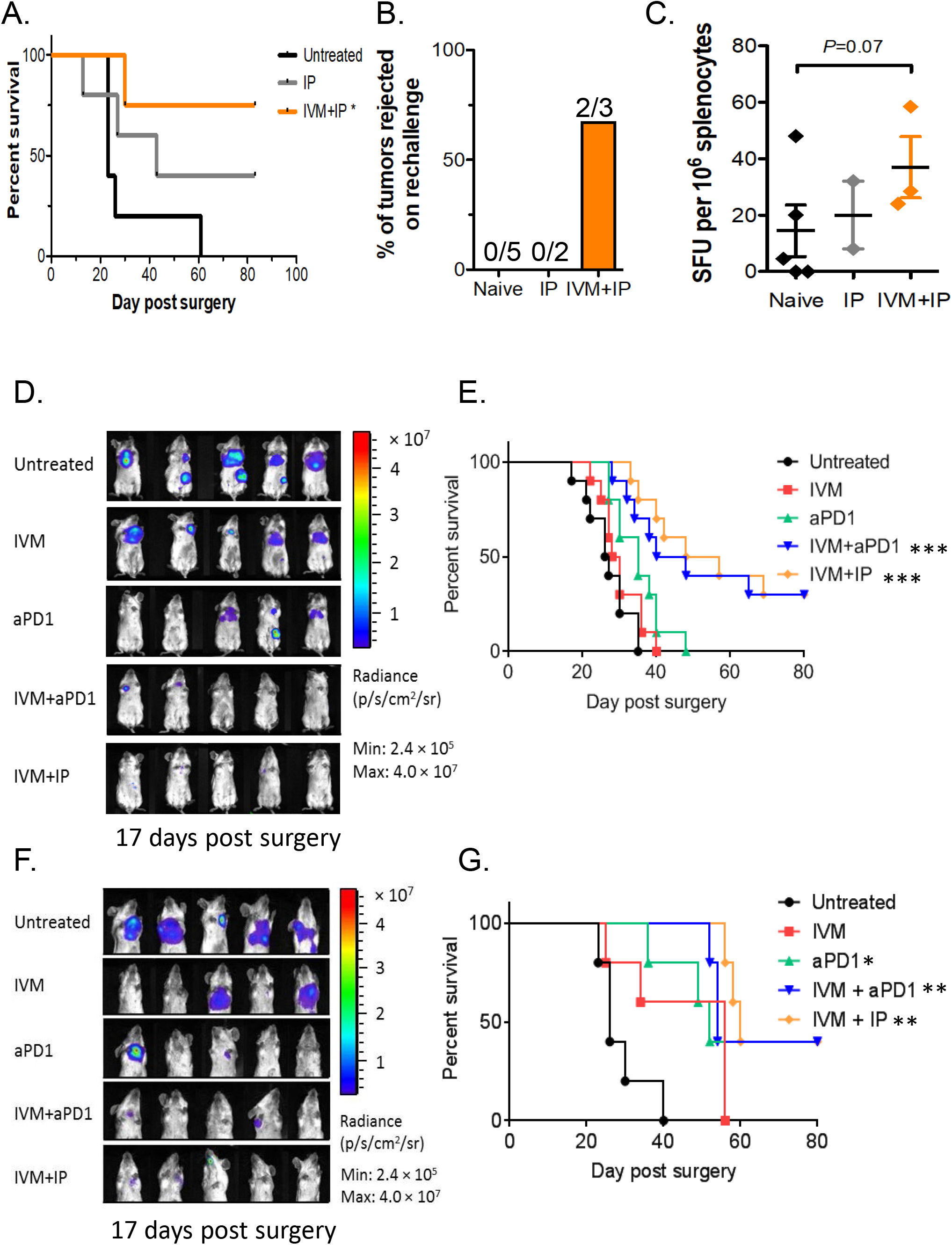
Combination ivermectin and IP therapy in the neoadjuvant, adjuvant, and metastatic settings. (A) Survival of animals following surgical resection of primary tumor (on day 16 post tumor inoculation). (B) Induction of protective immunity in treated mice that survived beyond Day 80, then re-challenged with 4T1 cells on the contralateral mammary fat pad. (C) IFNγ ELISPOT analysis of 4T1-reactive splenocytes in treated animals. Mean ± s.d., n = 5 mice, pooled data from 2 independent experiments. * p ≤ 0.05, ** p ≤ 0.01, *** p ≤ 0.001. (D) In vivo bioluminescence imaging of mice (on Day 17 post surgery and after completion of the entire treatment schedule) treated with ivermectin, anti-PD1, ivermectin + anti-PD1 +/−IL-2 (IP), or control in the adjuvant setting. Mean ± s.d., n = 5 mice, pooled data from 2 independent experiments. (E) Survival of animals in the adjuvant setting following surgical resection of primary tumor burden and treated starting 2 days after with ivermectin, anti-PD1, ivermectin + anti-PD1 +/−IL-2 (IP), or control. n = 5 mice per group, two-tailed log-rank test. ** p ≤ 0.01, **** p ≤ 0.0001. (F) In vivo bioluminescence imaging (on Day 17 post surgery and after completion of the entire treatment schedule) of mice with documented metastasis, then treated with ivermectin, anti-PD1, ivermectin + anti-PD1 +/−IL-2 (IP), or control. Mean ± s.d., n = 5 mice, pooled data from 2 independent experiments. (G) Kaplan-Meier survival analysis of mice in the metastatic setting treated with ivermectin, anti-PD1, ivermectin + anti-PD1 +/− IL-2 (IP), or control. n = 5 mice per group, two-tailed log-rank test. * p ≤ 0.05, ** p ≤ 0.01, *** p ≤ 0.001.

Surgery remains the primary treatment for breast cancer; however, relapse is common necessitating adjuvant therapy in high-risk patients post-surgery. We assessed the efficacy of ivermectin, anti-PD1, and recombinant IL-2 alone or in combination as adjuvant immunotherapy after surgery. 4T1 cells expressing luciferase (0.5×10^6^, 4T1-Luc) were injected into the mammary pad of female BALB/c mice and allowed to grow into palpable tumors over 10 days, after which tumors were surgically resected. Treatment was initiated on day 2 following surgery to mimic adjuvant therapy (schema in Fig. S1A). Development of recurrence and metastasis was monitored at multiple time points via bioluminescence imaging (Day 17 shown, Fig. 4D), then animals were monitored until they met euthanasia criteria based on decline in body condition score and signs of morbidity. Treatment with anti-PD1 or IVM alone led to similar survival as untreated controls (Fig. 4E). Animals treated with the combination of ivermectin and anti-PD1 (with or without IL-2) had significantly prolonged survival, with approximately 40% of animals becoming long-term survivors (p<0.001, Fig. 4E). Through statistical modeling, the ivermectin and anti-PD1 combination was found to be highly synergistic compared to IVM or anti-PD-1 alone (submodel p=0.007, FDR 2%, Table 1B). Interestingly, addition of IL-2 did not further enhance the survival benefit from the ivermectin and anti-PD1 combination (submodel p=0.51, FDR 67%, Table 1B). These data demonstrate that treatment with ivermectin and anti-PD1 (with or without IL-2) is effective in the adjuvant setting, without evidence for drug related or synergistic toxicity based on parallel body weight observations (Fig. S1B).

Metastasis is the main cause of death in cancer patients including breast cancer. To test the efficacy of this combination in the metastatic setting, we delayed treatment until at least 25% of animals post-surgery had detectable metastasis (generally day 7 after surgical resection of primary tumor). Progression of metastasis was monitored via bioluminescence imaging (schema in Fig. S1A), and animals were monitored until they met euthanasia criteria based on decline in body condition score and signs of morbidity (examples shown in Fig. 4F). All untreated animals required euthanasia due to metastatic disease around day 20-40 following surgical resection of primary tumor (Fig. 4G). Treatment with IVM alone led to modest, non-significant prolongation of survival as compared to untreated controls (Fig. 4G). Survival was slightly prolonged in animals treated with anti-PD1 only (p<0.05), but all animals required euthanasia by Day 60 as in the IVM alone group. Survival was significantly prolonged in animals treated with ivermectin and anti-PD1 (p<0.001), or ivermectin, anti-PD1 and IL-2 (p<0.01) as compared to untreated controls (Fig. 4G). Approximately 40% of animals on the combination therapy become long-term survivors. The combined effect of IVM and anti-PD-1 on survival in the metastatic setting was again found to be highly synergistic compared to IVM or anti-PD-1 alone (submodel p<0.001, FDR <1%, Table 1C). As in the adjuvant setting, addition of IL-2 did not further enhance the survival benefit from the ivermectin and anti-PD1 combination (submodel p=0.64, FDR 73%, Table 1C). These data demonstrate that treatment with ivermectin and anti-PD1 (with or without IL-2) is also effective in the metastatic setting.

## Discussion

Since its discovery in the mid-1970s, ivermectin has been used safely by over 700 million people worldwide to treat river blindness and lymphatic filariasis (23); it is inexpensive and accessible. Our results demonstrate that treatment with ivermectin induces robust T cell infiltration into breast tumors via induction of ICD, thus turning ‘cold’ tumors ‘hot’. Unlike conventional chemotherapy drugs, this agent has the added benefit of not suppressing host immune function, but rather has beneficial immunomodulatory effects – making it a promising and mechanistic partner for immune checkpoint blockade. The release and accumulation of high levels of extracellular ATP has emerged as a key characteristic feature of the tumor microenvironment (24), and a hallmark of ICD. We and others have previously shown that ivermectin is a positive allosteric modulator of purinergic signaling and the ATP/P2×4/P2×7/Pannexin-1 axis which operates in both cancer and immune cells (17, 25). In murine splenocytes treated *ex vivo*, we showed that ivermectin can selectively target various immune subpopulations in a P2×7-dependent fashion (Fig. 2B, C) and has immune-potentiating activities associated with augmented ratios of immune effectors versus immunosuppressive populations, including Tregs and myeloid cells (Fig. 2A, D). The observed selective targeting of different immune populations by ivermectin is consistent with previous reports demonstrating that mouse splenic Tregs (CD4+CD25+) have higher sensitivity to increasing (>100 μM) doses of extracellular ATP compared to CD8+ and CD4+CD25-T cells (26). This differential sensitivity to extracellular ATP is P2×7-dependent and directly associated with levels of surface P2×7 receptor expression (CD4+CD25+ > CD4+CD25-> CD8+ T cells). Recent reports showed that the ATP/P2×7 axis also operates in MDSC and MDSC-mediated immunosuppression (27, 28). This is consistent with our finding that ivermectin can selectively target expanded myeloid cells isolated from tumor-bearing mice *ex vivo* in a P2×7-dependent fashion. Further research will be needed to elucidate the relative sensitivities of different subsets of MDSC and tumor-associated macrophages/neutrophils (TAMs/TANs) to ivermectin, as well as to validate the *in vivo* effects of ivermectin on various myeloid subsets within the tumor microenvironment and systemically.

While differential ATP/P2×7-dependent cytotoxicity may be one possible explanation for the immunomodulatory effects of ivermectin *in vivo*, recent reports also implicate ATP release and P2×4-dependent signaling in the CXCL12/CXCR4-mediated migration and inflammation-driven recruitment of T cells (29). The role of P2×4 in T cell activation, proliferation and migration was particularly pronounced in CD4 T cells, which is consistent with our own data demonstrating ivermectin to be particularly potent at increasing the CD4+ Teff/Treg ratios in *ex vivo* treated splenocytes (Fig. 2D) and augmenting intra-tumoral infiltration with CD4+ T cells (Fig. 3D). Thus, infiltration of tumors by T cells in ivermectin treated mice may reflect a combination of selective depletion of suppressive cells as well as recruitment effects. The synergistic activity between ivermectin and anti-PD-1 checkpoint blockade at driving T cell infiltration into the tumor microenvironment is particularly intriguing as PD-1 functions as a negative feedback regulator of TCR signaling. P2×4/P2×7-gated Pannexin-1(PANX1) opening and ATP release play a central role in T cell activation by providing a feed-forward loop for TCR-initiated and ATP-driven ATP release at the immunological synapse. The ability of ivermectin as an allosteric modulator of P2×4/P2×7/PANX1 receptors to modulate purinergic signaling operating in both cancer and immune cells therefore may be enhanced by elevated levels of ATP within the tumor microenvironment and the immunological context, including magnitude of chemokine/TCR signaling and chemokine/TCR-driven ATP release. Consistent with the latter possibility, we demonstrated that the potentiating effect of ivermectin on the Teff/Treg ratio appears to be stronger and sustained in the presence of TCR stimulation (Fig. 2D). Further studies will be needed to unravel how these multi-faceted effects of ivermectin to induce immunogenic cancer cell death, differentially modulate immune cells, and harness the ATP-rich tumor microenvironment may all contribute to its ability to synergize with immune checkpoint blockade *in vivo*.

Immune checkpoint inhibitors (ICI) are effective as single agents only in a small subset of cancer patients. Hundreds of clinical trials are currently testing various combinations of ICI with FDA-approved or experimental agents. Such combinations are mainly put together based on partial efficacy of the partnering agent with little or no mechanistic rationale for synergy. Importantly, recent analyses found no evidence from any trial data reported to date that ICIs are synergistic or additive with other drugs (30), but instead synergistic toxicity may be observed (31, 32). We showed that ivermectin represents a rational mechanistic partner for immune checkpoint blockade, demonstrating bona fide synergy when neither agent worked alone. Synergy between PD-1 blockade and ivermectin is mechanistically associated with the ability of the ivermectin to drive immunogenic cancer cell death and T cell infiltration into tumors, thus converting ‘cold’ tumors ‘hot’ (33). This combination led to complete resolution of the primary tumor in a significant fraction of animals, and with protective anti-tumor immunity in the responders. We went on to demonstrate that this novel combination is effective in the neoadjuvant, adjuvant, and metastatic settings that mimic clinical situations in which it may be used. Based on its novel dual mechanisms of action in cancer, ivermectin may also potentiate the anti-tumor activity of other FDA-approved immune checkpoint inhibitors. Lastly, ivermectin is inexpensive, making it attainable for everyone including cancer patients in developing countries. The preclinical findings we present suggest that the combination of ivermectin and anti-PD1 antibody merits clinical testing in breast cancer patients.

## Acknowledgments

Authors wish to acknowledge Dr. Steve Vonderfecht for advice and help on animal studies; Dr. Larry Wong and Gilbert Acosta for technical help; and Dr. Chris Gandhi for critical review of this manuscript.

## Author contributions

Conceptualization: PPL, DD, DJI, NB; Experimental work: DD, ZH, NB; Data analysis: DD, ZH, NB; Writing: DD, ZH, PPL, DJI, NB

## Data and materials availability

Data required to reproduce the statistical analyses reported in the paper will be deposited in the Supplementary Materials. Access to those data is unrestricted.

## Materials and methods

### Mice and treatment

Female BALB/c mice were purchased from Charles River Laboratories at 5-8 weeks of age and housed in City of Hope’s animal care facilities under pathogen-free conditions. All procedures were performed under approval from City of Hope’s Animal Care and Use Committee. Mice were inoculated with 100,000 4T1 breast cancer cells in the right mammary fat pad, then palpated to check for tumor engraftment before commencing their assigned treatment regimen. Treatments included: 5 mg/kg of ivermectin (Sigma Aldrich, St. Louis MO) given via oral gavage daily for 6 days; 10 mg/kg of anti-PD1 (BioXCell, West Lebanon NH) treatment given subcutaneously once weekly; MSA-IL2 administered at 1.5 mg/kg by intraperitoneal injection in 50 µL sterile PBS once weekly; combination treatments; or no treatment (Fig. S1). Ivermectin was solubilized in 45% (2-Hydroxypropyl)-β-cyclodextrin (Sigma Aldrich, 332593-1KG), as previously described (34). Tumor growth was measured 2-3 times a week with a digital caliper for up to 56 days. Mice were euthanized when tumor growth reached 1.5 cm in length or width. Tumor volume was calculated as (length*width^2^)/2. Metastasis experiments were performed by injecting 0.5×10^6^ luciferase expressing 4T1 tumor cells (4T1-Luc) subcutaneously in the mammary gland of female BALB/c mice, followed by surgically resection of the primary tumor on day 14 after inoculation. In vivo bioluminescence imaging was used to monitor metastatic outgrowth, which was carried out on a Lago X optical imaging system (Spectral Instruments Imaging, Tucson, AZ). Overall tumor burden per mouse was assessed weekly via bioluminescence imaging. Recurrence of primary tumor was recognized when the animal’s luciferase value exceeded 600,000 photons/sec/cm^2^/steradian, a threshold chosen because it was well above the lower limit of reproducible detection (510,000) and because, in optimization experiments, 600,000 was the lowest threshold consistently followed by ever-increasing values and eventually death. There was no significant toxicity following treatment with oral ivermectin combined with systemic anti-PD1 and IL-2 as measured by weight loss (Fig. S1).

### Neoadjuvant Setting Mice and Treatment

Female BALB/c mice were purchased from The Jackson Laboratories at 6-8 weeks of age and maintained in animal care facilities under pathogen-free conditions at the Massachusetts Institute of Technology. All procedures were performed under approval from MIT’s Animal Care and Use Committee.

An inoculum of 0.5×10^6^ 4T1-Luc tumor cells were injected subcutaneously (s.c.) in the mammary gland in 100 µL sterile PBS. Tumor onset was monitored by palpation (usually 3-5 days after inoculation). Six days following inoculation, mice were randomized into treatment groups and treatment was performed as indicated in Supp. Fig. 2. A dose of 5 mg/kg ivermectin was administered by oral gavage in 50 µL sterile PBS. Anti-PD-1 (clone RMP1-14, BioXCell) was administered at 10 mg/kg by intraperitoneal injection in 50 µL sterile PBS. MSA-IL2 was administered at 1.5 mg/kg by intraperitoneal injection in 50 µL sterile PBS.

Surgical resection of primary tumor was performed on day 16 following tumor inoculation. Mice were injected with the analgesic sustained-release Buprenorphine (ZooPharm, 1 mg/kg body weight) and meloxicam (2 mg/kg body weight) by subcutaneous injection. Animals were anesthetized with isoflurane and complete anesthetization was confirmed by lack of a toe pinch reflex. The surgical area was shaved and sterilized by swabbing with alternating application of betadine surgical scrub and 70% ethanol. The tumor and surrounding mammary fat pad was removed by blunt dissection using autoclaved surgical instruments (Braintree Scientific). Wounds were closed using 4-0 nylon monofilament sutures with a 3/8 reverse cutting needle (Ethilon). Mice were monitored for consciousness in a warm, dry area immediately post-operation. Thereafter, mice were dosed with meloxicam (2 mg/kg body weight) at 24 hour intervals for 3 days post-surgery. Sutures were removed at 7-10 days post-operation.

Mice were monitored for development of metastasis starting at day 10-14 following surgical resection of the primary tumor. Animals were injected i.p. with sterile-filtered D-luciferin (Xenogen) in PBS (150 mg/kg body weight in 200 µL) and anesthetized with isoflurane. Bioluminescence images were collected at 10 minutes following injection with a IVIS Spectrum Imaging System (Xenogen). Acquisition times ranged 10-30 seconds. Images were analyzed using Living Image software (Xenogen). Animals were monitored daily for morbidity and euthanized if signs of distress were observed, including but not limited to difficulty in ambulating or breathing, significant weight loss (>20% starting body weight), poor body condition (score <2) or veterinary staff recommendation. Necropsy was performed to confirm presence of visible metastatic nodules.

To evaluate response to re-challenge, mice that survived metastasis development following surgical resection or naïve control mice were challenged with a subcutaneous injection of 0.1×10^6^ 4T1-Luc cells in 100 µL sterile PBS in the flank opposite the site of the primary tumor. The mice were subsequently monitored every 2-3 days for tumor growth at the inoculation site.

### ELISPOT assay

Target 4T1-Luc cells were treated with mouse IFN-gamma (Peprotech) for 18 hours, washed, and irradiated (120 Gy). Splenocytes were isolated from untreated or treated mice on day 16 following to tumor re-challenge. Quantification of IFN-g response was determined using a BD mouse IFN-g ELISPOT kit. Target cells were seeded at 0.025×10^6^ cells per well. Effector cells were seeded 1.0×10^6^ cells per well. Plates were wrapped in foil and incubated at 37°C for 24 hours and developed following the manufacture’s protocol. Plates were scanned using a CTL-ImmunoSpot plate reader and spots were enumerated using CTL ImmunoSpot software.

### Tissue staining and quantification

Tumors were isolated from mice and sectioned into 5 micrometer sections for staining with the desired markers (below) using Tyramide Signal Amplification (PerkinElmer, Waltham MA) per manufacturer’s protocol. Whole tumor images were scanned using the Vectra 3 Automated imaging system (PerkinElmer) and quantified using the ImagePro analysis software.

### Flow cytometry

Cell surface markers were stained for 30 minutes in the dark at 4°C. Intracellular cytokine staining was performed using the ebioscience Foxp3 staining kit (Thermo Fisher Scientific, Waltham MA) per manufacturer’s protocol. The following mouse antibodies from BioLegend (San Diego CA) were used: CD4 (GK1.5); CD8 (53-6.7); Tbet (4B10); Gata3 (16E10A23); Foxp3 (MF-14); IFNγ (XMG1.2); IL-10 (JES5-16E3); IL17 (TC11-18H10.1); and TGFβ (TW7-16B4). RORγt (AFKJS-9) was ordered from eBioscience (ThermoFisher Scientific). To show T cell reactivity, splenocytes were isolated from tumor bearing mice and cultured with 4T1 cells at a ratio of 5:1 (splenocytes to tumor cells) in the presence anti-CD107A/CD107B (ThermoFisher Scientific) and Monensin for 4 hours. After 4 hours, cells were stained for surface and intracellular markers described above. Flow cytometry analysis of T cell markers on human PBMCs was performed using the following clones: CD8 (RPA-T8); CD4 (SK3); Tbet (4B10); Ki67 (Ki67) from BioLegend; RORγt and granzyme B (GB11) from ThermoFisher Scientific.

### Statistical analysis

Mean values were compared using t tests. Data on tumor volume over time were log-transformed prior to statistical modeling; prior to transformation, values of zero were replaced with 0.1. complete response (CR) to treatment was defined as permanent shrinkage of tumor volume to zero at any time during follow-up; no tumor that shrank to zero resumed growth. The competing survival outcome was progression, defined as tumor growth beyond 150 mm^3^, after which tumors never underwent CR but instead became necrotic or large, necessitating euthanasia. The follow-up of subjects that experienced neither CR nor progression was censored at last observation, except when the last available tumor measurement fell just short of the 150 mm^3^ threshold for progression; in such cases (n=2, volume 139 and 141 mm^3^, respectively, at final measurement on Day 25), progression was assumed to occur by what would have been the next scheduled measurement.

Cumulative incidence of competing outcomes was calculated and plotted according to Gray(35). The related outcomes of tumor volume, CR, and progression were modeled jointly(36). The submodel of longitudinal tumor volume used linear mixed regression, while the survival submodels of CR and progression used parametric hazard regression with Weibull function. Statistical significance was defined as p<0.05. A greater-than-additive (synergy) effect of combination therapy was demonstrated when the sum of effects of each drug alone fell outside the 95% confidence interval around the effect of combination therapy. To maximize statistical power and obtain unbiased results despite the non-random missingness of longitudinal data due to death, each pair of outcome measures per trial was modeled jointly (36). Each joint model included a linear mixed effects submodel of the longitudinal outcome and a survival submodel. To keep the trials’ overall risk of error below 5%, p values for the primary hypothesis for synergy from combination treatment were subjected to the step-up Bonferroni adjustment of Hochberg (37). Separately, p values for the secondary hypotheses underwent the same adjustment.

## Supplementary Materials

**Supplementary Table 1.**
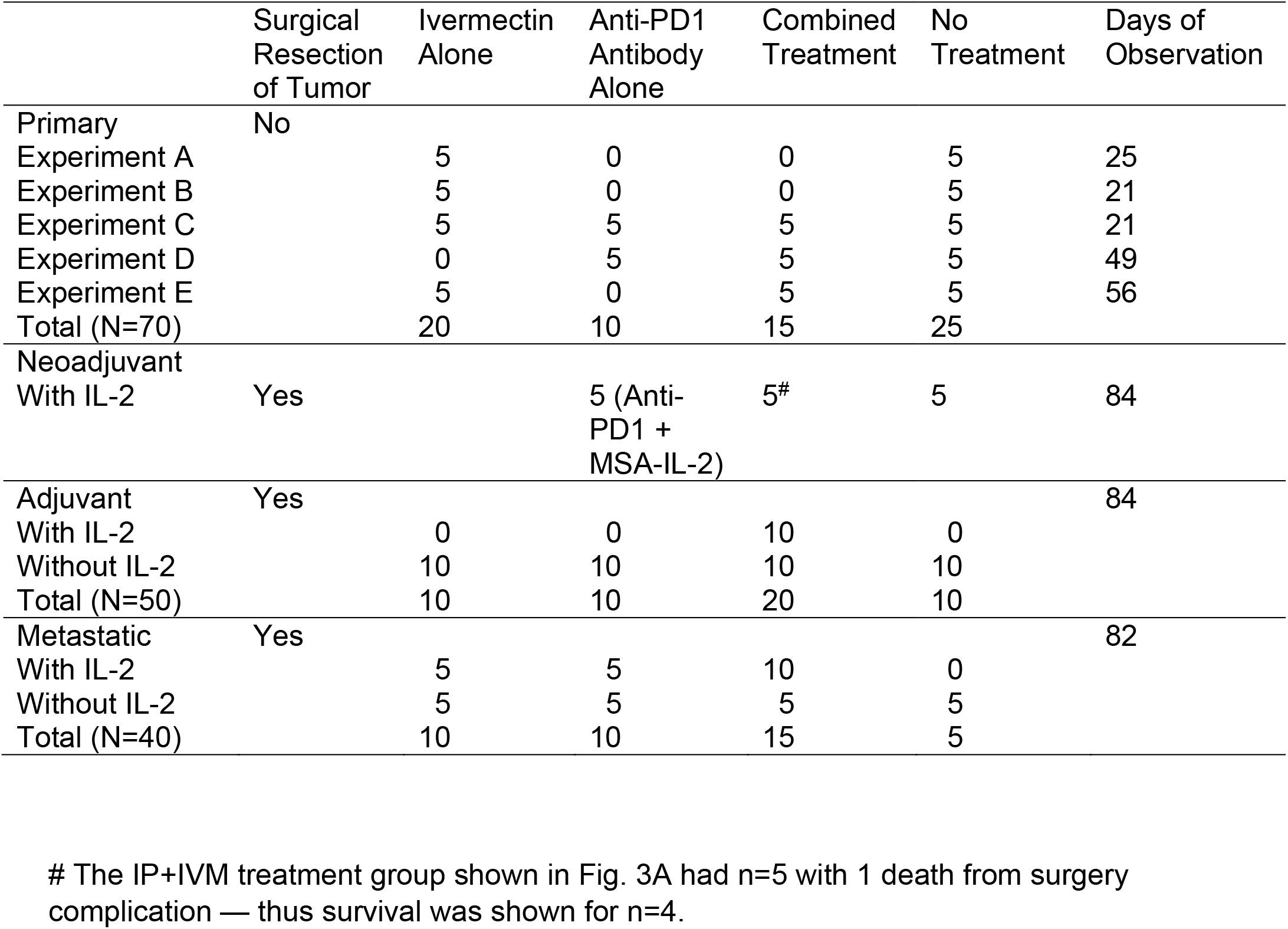
Experimental Design, Treatment Settings.

**Fig S1.**
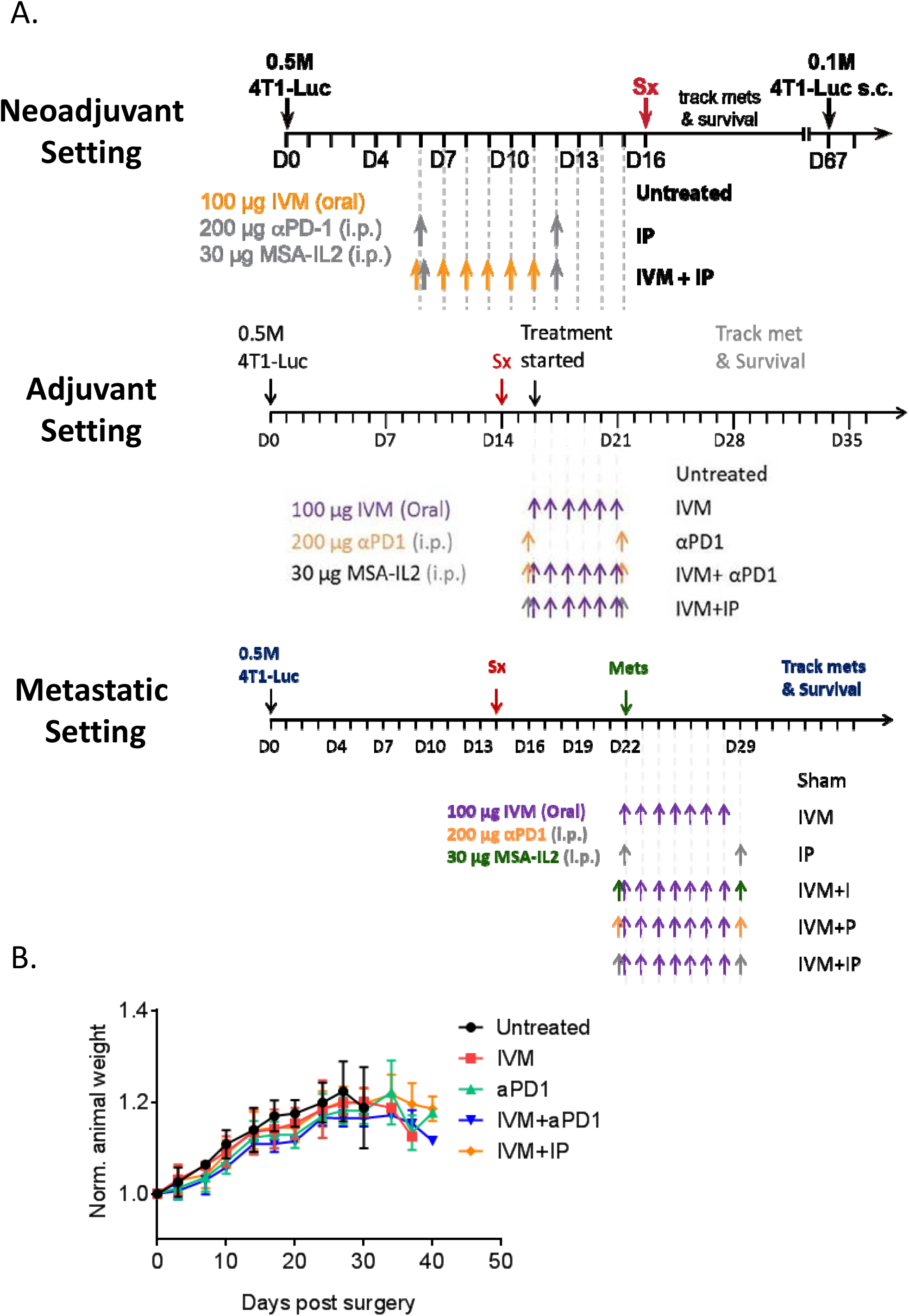
(A) Treatment schedules in the neoadjuvant, adjuvant, and metastatic settings (surgical resection = **Sx**). (B) Body weight measurements of treated animals demonstrating the absence of significant synergistic toxicity associated with the combination of anti-PD-1 and Ivermectin in the adjuvant settings. Similar observations were seen in the metastatic treatment settings.

**Fig S2.**
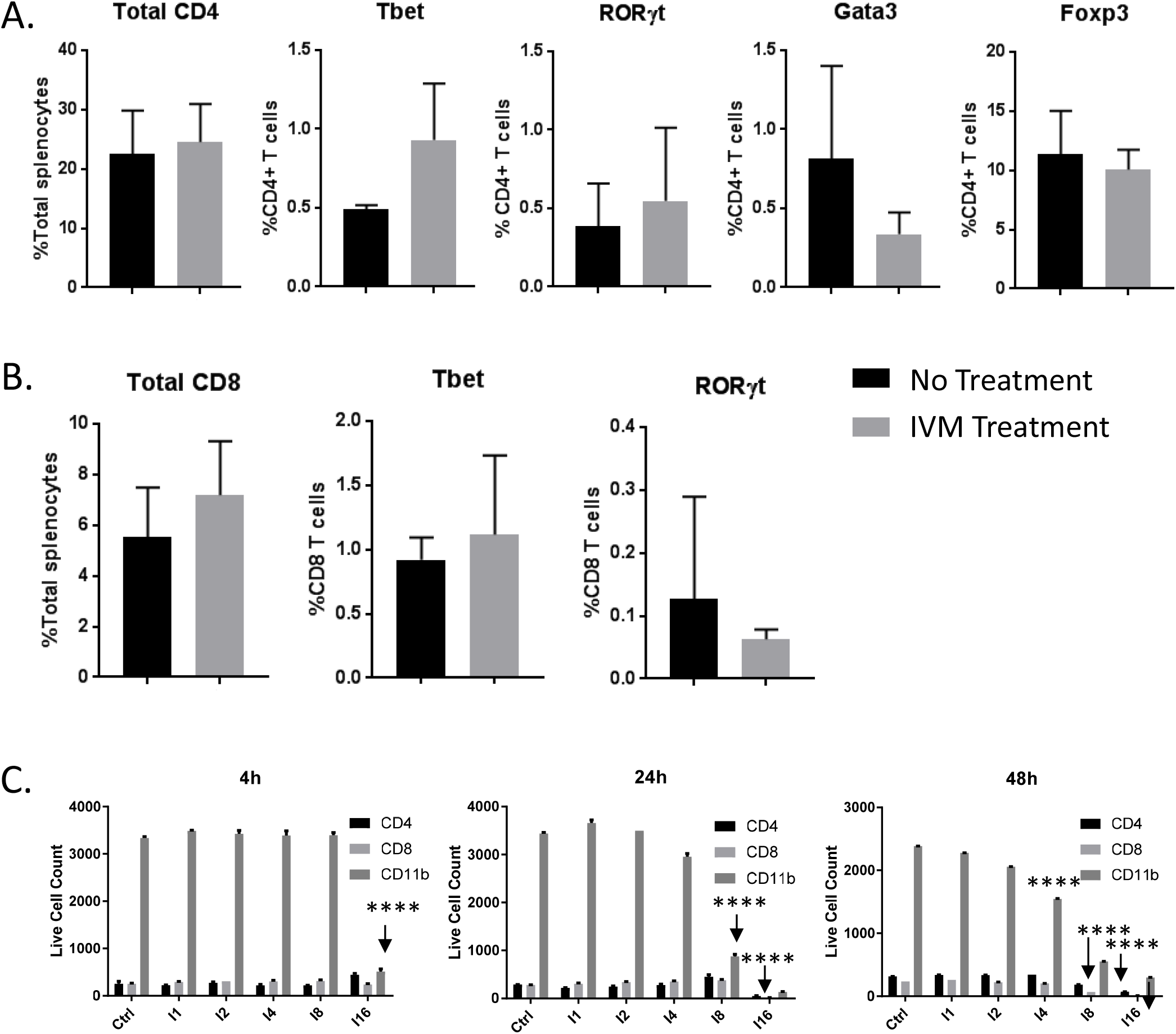
Immunomodulatory effects of Ivermectin on immune cells *in vivo* and *ex vivo*. (A) Flow cytometry analysis of splenocytes from 4T1 tumor-bearing animals treated with ivermectin, demonstrating the absence of significant changes *in vivo* of various CD4 (A) and CD8 (B) effector and regulatory T cell subpopulations, which were identified based on the expression of key transcriptional factors as indicated. All comparisons were non-significant, NS. (C) Differential sensitivity of immune subpopulations in splenocytes isolated from 4T1 tumor-bearing mice exposed *ex vivo* to increasing (1-16 μM) doses of ivermectin for 4h to 48h showing dose and time-dependent sensitivity. A linear mixed effects model of log cell count adjusted for cell type revealed that the CD11b+ myeloid cells were the most sensitive to ivermectin, showing significant reductions with as little as 4 μM after 48 hours, or 8 μM after 24 hours, or 16 μM after 4 hours (each result, p<0.0001). In contrast, achieving similar reductions in lymphocytes required higher doses and/or longer exposure to ivermectin, being observed in CD8+ cells only after 48 hours of 8 μM or 24 hours of 16 μM and in CD4+ cells only after the maximum exposure (48 hours of 16 μM). Statistical significance versus (-) CTRL or as indicated was evaluated using the linear mixed effects model of log cell count adjusted for cell type, **** p ≤ 0.0001.

## References

1. Pardoll DM. The blockade of immune checkpoints in cancer immunotherapy. Nature reviews Cancer. 2012;12(4):252–64.

2. Sharma P, and Allison JP. The future of immune checkpoint therapy. Science. 2015;348(6230):56–61.

3. Disis ML, and Stanton SE. Triple-negative breast cancer: immune modulation as the new treatment paradigm. American Society of Clinical Oncology educational book American Society of Clinical Oncology Meeting. 2015:e25–30.

4. Nanda R, Chow LQ, Dees EC, Berger R, Gupta S, Geva R, et al. Pembrolizumab in Patients With Advanced Triple-Negative Breast Cancer: Phase Ib KEYNOTE-012 Study. Journal of clinical oncology : official journal of the American Society of Clinical Oncology. 2016;34(21):2460–7.

5. Galluzzi L, Buque A, Kepp O, Zitvogel L, and Kroemer G. Immunogenic cell death in cancer and infectious disease. Nature reviews Immunology. 2017;17(2):97–111.

6. Kroemer G, Galluzzi L, Kepp O, and Zitvogel L. Immunogenic cell death in cancer therapy. Annual review of immunology. 2013;31:51–72.

7. Galluzzi L, Vitale I, Warren S, Adjemian S, Agostinis P, Martinez AB, et al. Consensus guidelines for the definition, detection and interpretation of immunogenic cell death. Journal for ImmunoTherapy of Cancer. 2020;8(1):e000337.

8. Apetoh L, Ghiringhelli F, Tesniere A, Obeid M, Ortiz C, Criollo A, et al. Toll-like receptor 4-dependent contribution of the immune system to anticancer chemotherapy and radiotherapy. Nature medicine. 2007;13(9):1050–9.

9. Michaud M, Martins I, Sukkurwala AQ, Adjemian S, Ma Y, Pellegatti P, et al. Autophagy-dependent anticancer immune responses induced by chemotherapeutic agents in mice. Science. 2011;334(6062):1573–7.

10. Casares N, Pequignot MO, Tesniere A, Ghiringhelli F, Roux S, Chaput N, et al. Caspase-dependent immunogenicity of doxorubicin-induced tumor cell death. The Journal of experimental medicine. 2005;202(12):1691–701.

11. Mattarollo SR, Loi S, Duret H, Ma Y, Zitvogel L, and Smyth MJ. Pivotal role of innate and adaptive immunity in anthracycline chemotherapy of established tumors. Cancer research. 2011;71(14):4809–20.

12. DeNardo DG, Brennan DJ, Rexhepaj E, Ruffell B, Shiao SL, Madden SF, et al. Leukocyte complexity predicts breast cancer survival and functionally regulates response to chemotherapy. Cancer discovery. 2011;1(1):54–67.

13. Denkert C, Loibl S, Noske A, Roller M, Muller BM, Komor M, et al. Tumor-associated lymphocytes as an independent predictor of response to neoadjuvant chemotherapy in breast cancer. Journal of clinical oncology : official journal of the American Society of Clinical Oncology. 2010;28(1):105–13.

14. Ladoire S, Mignot G, Dabakuyo S, Arnould L, Apetoh L, Rebe C, et al. In situ immune response after neoadjuvant chemotherapy for breast cancer predicts survival. The Journal of pathology. 2011;224(3):389–400.

15. Halama N, Michel S, Kloor M, Zoernig I, Benner A, Spille A, et al. Localization and density of immune cells in the invasive margin of human colorectal cancer liver metastases are prognostic for response to chemotherapy. Cancer research. 2011;71(17):5670–7.

16. Ray-Coquard I, Cropet C, Van Glabbeke M, Sebban C, Le Cesne A, Judson I, et al. Lymphopenia as a prognostic factor for overall survival in advanced carcinomas, sarcomas, and lymphomas. Cancer research. 2009;69(13):5383–91.

17. Draganov D, Gopalakrishna-Pillai S, Chen YR, Zuckerman N, Moeller S, Wang C, et al. Modulation of P2×4/P2×7/Pannexin-1 sensitivity to extracellular ATP via Ivermectin induces a non-apoptotic and inflammatory form of cancer cell death. Scientific reports. 2015;5:16222.

18. Kepp O, Senovilla L, Vitale I, Vacchelli E, Adjemian S, Agostinis P, et al. Consensus guidelines for the detection of immunogenic cell death. Oncoimmunology. 2014;3(9):e955691.

19. Rubio V, Stuge TB, Singh N, Betts MR, Weber JS, Roederer M, et al. Ex vivo identification, isolation and analysis of tumor-cytolytic T cells. Nature medicine. 2003;9(11):1377–82.

20. Boyman O, Krieg C, Letourneau S, Webster K, Surh CD, and Sprent J. Selectively expanding subsets of T cells in mice by injection of interleukin-2/antibody complexes: implications for transplantation tolerance. Transplantation proceedings. 2012;44(4):1032–4.

21. Moynihan KD, Opel CF, Szeto GL, Tzeng A, Zhu EF, Engreitz JM, et al. Eradication of large established tumors in mice by combination immunotherapy that engages innate and adaptive immune responses. Nature medicine. 2016;22(12):1402–10.

22. Zhu EF, Gai SA, Opel CF, Kwan BH, Surana R, Mihm MC, et al. Synergistic innate and adaptive immune response to combination immunotherapy with anti-tumor antigen antibodies and extended serum half-life IL-2. Cancer cell. 2015;27(4):489–501.

23. Crump A, and Omura S. Ivermectin, ‘wonder drug’ from Japan: the human use perspective. Proceedings of the Japan Academy Series B, Physical and biological sciences. 2011;87(2):13–28.

24. Burnstock G, and Di Virgilio F. Purinergic signalling and cancer. Purinergic signalling. 2013;9(4):491–540.

25. Junger WG. Immune cell regulation by autocrine purinergic signalling. Nature reviews Immunology. 2011;11(3):201–12.

26. Aswad F, and Dennert G. P2×7 receptor expression levels determine lethal effects of a purine based danger signal in T lymphocytes. Cellular immunology. 2006;243(1):58–65.

27. Bianchi G, Vuerich M, Pellegatti P, Marimpietri D, Emionite L, Marigo I, et al. ATP/P2×7 axis modulates myeloid-derived suppressor cell functions in neuroblastoma microenvironment. Cell Death Dis. 2014;5:e1135.

28. Principi E, and Raffaghello L. The role of the P2×7 receptor in myeloid-derived suppressor cells and immunosuppression. Curr Opin Pharmacol. 2019;47:82–9.

29. Ledderose C, Liu K, Kondo Y, Slubowski CJ, Dertnig T, Denicolo S, et al. Purinergic P2×4 receptors and mitochondrial ATP production regulate T cell migration. The Journal of clinical investigation. 2018;128(8):3583–94.

30. Palmer AC, Izar B, and Sorger PK. Combinatorial benefit without synergy in recent clinical trials of immune checkpoint inhibitors. medRxiv. 2020:2020.01.31.20019604.

31. Boutros C, Tarhini A, Routier E, Lambotte O, Ladurie FL, Carbonnel F, et al. Safety profiles of anti-CTLA-4 and anti-PD-1 antibodies alone and in combination. Nat Rev Clin Oncol. 2016;13(8):473–86.

32. Gao L, Yang X, Yi C, and Zhu H. Adverse Events of Concurrent Immune Checkpoint Inhibitors and Antiangiogenic Agents: A Systematic Review. Front Pharmacol. 2019;10:1173.

33. Duan Q, Zhang H, Zheng J, and Zhang L. Turning Cold into Hot: Firing up the Tumor Microenvironment. Trends Cancer. 2020;6(7):605–18.

34. Melotti A, Mas C, Kuciak M, Lorente-Trigos A, Borges I, and Ruiz i Altaba A. The river blindness drug Ivermectin and related macrocyclic lactones inhibit WNT-TCF pathway responses in human cancer. EMBO molecular medicine. 2014;6(10):1263–78.

35. Gray R. cmprsk: Subdistribution Analysis of Competing Risks. R package version 2.2-7. https://CRAN.R-project.org/package=cmprsk

36. Wang W, Wang W, Mosley TH, and Griswold ME. A SAS macro for the joint modeling of longitudinal outcomes and multiple competing risk dropouts. Computer methods and programs in biomedicine. 2017;138:23–30.

37. Hochberg Y. A Sharper Bonferroni Procedure for Multiple Tests of Significance. Biometrika. 1988;75(4):800–2.

